# Functional genetic characterization by CRISPR-Cas9 of two enhancers of *FOXP2* in a child with speech and language impairment

**DOI:** 10.1101/064196

**Authors:** R. Torres-Ruiz, A. Benitez-Burraco, M. Martínez-Lage, S. Rodríguez-Perales, P. García-Bellido

**Author notes:** Corresponding authors: Sandra Rodriguez-Perales. Paloma Garcia-Bellido.

## Abstract

Mutations in the coding region of *FOXP2* are known to cause speech and language impairment. Microdeletions involving the region downstream the gene have been also associated to speech and cognitive deficits. We recently described a girl harbouring a complex chromosomal rearrangement with one breakpoint downstream the gene that might affect her speech and cognitive abilities via physical separation of distant regulatory DNA elements. In this study, we have used highly efficient targeted chromosomal deletions induced by the CRISPR/Cas9 genome editing tool to demonstrate the functionality of two enhancers, FOXP2-E^proximal^ and FOXP2-E^distal^, located in the intergenic region between *FOXP2* and its adjacent *MDFIC* gene. Deletion of any of these two functional enhancers in the neuroblastomic cell line SK-N-MC downregulates *FOXP2* and decreases FOXP2 protein levels, conversely it upregulates *MDFIC* and increases MDFIC protein levels. This suggests that both regulatory elements may be shared between *FOXP2* and *MDFIC*. We expect these findings contribute to a deeper understanding of how *FOXP2* and *MDFIC* are regulated to pace neuronal development supporting speech and language.

## INTRODUCTION

Mutations in the gene *FOXP2*, encoding a transcription factor, are known to cause speech and language impairment (Vargha-Khadem et al. 2005; Zhao et al. 2010). Polymorphisms of the gene have been also associated to schizophrenia (Tolosa et al. 2010) and frontotemporal lobar degeneration (Padovani et al. 2010). *FOXP2* has been hypothesised to regulate the development and function of brain areas involved in human language processing (Lai et al., 2003; Fisher and Scharff 2009), because of its known role in neurogenesis, neuron differentiation and migration patterns in the developing telencephalon in mice (Tsui et al. 2013; Chiu et al. 2014; Garcia-Calero et al. 2016). Pathogenic mutations in humans have proven to impair auditory-motor association learning in mice (Kurt et al. 2012). Nonetheless, the exact role of *FOXP2* in normal development is unknown. Common variants of the gene do not contribute appreciably to individual differences in language development (Mueller et al. 2016), nor in brain structure (Hoogman et al. 2014), although a *FOXP2* polymorphism has been associated with enhanced procedural learning of non-native speech sound categories (Chandrasekaran et al. 2015). Less is known about how the expression of the gene is modulated. The promoter of *FOXP2* contains four transcription start sites (Schroeder and Myers 2008), with multiple alternative splicing sites (Bruce and Margolis 2002). *FOXP2* also contains six ultraconserved regions in its introns (Bejerano et al. 2004, Schroeder and Myers 2008), as well as six predicted enhancers for Lef1 (Hallikas et al 2006). Lef1 is a transcription factor that drives expression of the *Foxp2* gene in the central nervous system during zebrafish embryogenesis (Bonkowsky et al. 2008). Interestingly, several microRNAS bind the 3’UTR of the gene and regulate the expression of *FOXP2* (Clovis et al. 2012; Shi et al. 2013; Fu et al. 2014a; Cuiffo et al. 2014).

Apart from gene mutations, microdeletions involving *FOXP2* and/or *MDFIC*, the adjacent gene downstream *FOXP2*, and the region between these two genes have been found in subjects with speech delay and cognitive impairment (DECIPHER patients 262086, 292652, and 301696). We have recently reported on a young female harbouring a genomic complex rearrangement involving chromosomes 7 and 11, who presents with severe expressive and receptive speech and language impairment in both Castilian Spanish and Valencian (Moralli et al. 2015). Although the *FOXP2* coding region is intact, the breakpoint in 7q31.1 is located 205.5 kb downstream the 3’ end of *FOXP2* and 22.8 kb upstream the 5’ region of *MDFIC*. Becker et al. (2015) found and characterized a functional enhancer located 2.5 kb downstream the breakpoint and hypothesized that rearrangement of this enhancer by chromosomal translocation may contribute to the observed language phenotype by disturbing *FOXP2* gene expression. In our proband this element was hypothesized to be in derivative chromosome 7q separated from *FOXP2.* While *FOXP2* was shown to have been rearranged to derivative chromosome 11p, *MDFIC* had been located in derivative chromosome 7q, based on the FISH results from the RP11-243D16 BAC (Moralli et al 2015). A more robust approach, aimed at looking for changes in the expression levels of the gene seems desirable in order to know if this enhancer regulates *FOXP2* expression.

The development of nuclease mediated genome editing tools, specially, of those based on clustering regularly interspaced short palindromic repeats (CRISPR) (Sakuma and Woltjen 2014; Torres-Ruiz and Rodriguez-Perales 2016), has emerged as a highly efficient way of inducing targeted chromosomal deletions and an accurate method to validate the functionality of enhancers (Cong et al. 2013; Mali et al. 2013). Here we report a detailed study of the intergenic region between the *FOXP2* and *MDFIC* genes. We have found that this region contains, apart from the enhancer reported in Becker et al. (2015), a second functional enhancer, FOXP2-E ^proximal^. We performed targeted deletions of each regulatory element by CRISPR-Cas9 and found that both affect the expression levels of *FOXP2* and *MDFIC* in an opposite manner, increasing FOXP2 and reducing MDFIC mRNA and protein levels. We hypothesise therefore that the breakpoint in this case would cause *FOXP2* to be anomalously downregulated by the separation of FOXP2-^distal^ from *FOXP2*, while *MDFIC* to be anomalously upregulated by the separation of FOXP2-^proximal^ from *MDFIC*. These changes in the expression levels of these two genes may account for the observed language deficits in this case. We expect these findings contribute to a better understanding of how *FOXP2* is regulated.

## MATERIALS AND METHODS

### Cell culture and electroporation

Cells of the human non neuronal cell-line HEK293A (CRL-1573, ATCC, USA) and the neuroblastomic cell-line SK-N-MC (HTB-10, ATCC, USA) were maintained under standard conditions in Dulbecco’s modified Eagle’s medium (DMEM) (Lonza), supplemented with 10% foetal bovine serum (FBS) (Life Technologies), 1% GlutaMAX (Life Technologies), and 10mg/ml penicillin/streptomycin (Life Technologies). The neuroblstomic cell-line SK-SY-5Y (CRL-2266, ATCC, USA) was cultured in a 1:1 mixture of Dulbecco’s modified Eagle’s medium (DMEM) (Lonza) and F12 medium (Lonza) supplemented with 10% FBS (Life Technologies), 1% GlutaMAX (Life Technologies), and 10mg/ml penicillin/streptomycin (Life Technologies). Cells were cultured at 37°C in a humidified incubator in a 5% CO_2_ + 20% O_2_ atmosphere.

For electroporation, we used the Neon Transfection System (Life Technologies). The manufacturer’s protocols for HEK293A, SK-N-MC and SH-SY5Y cells were modified as follows. The three cell types were electroporated at 80% confluence. Cells were trypsinized and resuspended in R solution (Life Technologies). For SK-N-MC and SH-SY5Y, 10-µl tips were used to electroporate 2.5×10^6^ cells with a single 50-ms pulse of 900 V. For HEK293A cells, 4×10^5^ cells were electroporated with 10-µl tips using three 10-ms pulses of 1245 V. After electroporation, cells were seeded in a 24-well plate containing pre-warmed medium. When required, cells were sorted 72 h post-transfection.

### Construction of Double-Guide Cas9-Encoding Plasmids

The parental pLV-U6^#1^H1^#2^-C9G vector has been described elsewhere (Torres et al. 2014a). Two gBlocks gene fragments were synthesized to clone sgRNA#1 and sgRNA#2 flanking the FOXP2-E^proximal^ and FOXP2-E^distal^ enhancer regions in the backbone vector using BsrGI and SpeI target sites.

### Flow Cytometry and Cell Sorting

72 hours after electroporation, cells were trypsinized and washed with DPBS twice, counted, and resuspended in an appropriate volume of sorting buffer (PBS containing 1% FBS and antibiotics) for flow cytometry analysis. Immediately before cell sorting, samples were filtered through a 70-µm filter to remove any clumps or aggregates. Cell sorting was carried out in a Synergy 2L instrument (Sony Biotechnology Inc.); flow cytometry was performed in a BD LSR Fortessa analyzer (BD Biosciences). Cells were sorted and seeded individually per well in a 96 well-plate.

### Genomic DNA Extraction and PCR Analysis

Genomic DNA was extracted using standard procedures (Torres et al. 2014b). Briefly, 5-10×10^6^ cells were either trypsinized or scraped, washed in PBS, pelleted, and lysed in 100mM NaCl, Tris (pH 8.0) 50mM, EDTA 100mM, and 1% SDS. After overnight digestion at 56°C with 0.5 mg/ml of proteinase K (Roche Diagnostics), the DNA was cleaned by precipitation with saturated NaCl, and the clear supernatant was precipitated with isopropanol, washed with 70% ethanol, air-dried, and resuspended overnight at room temperature in 1xTE buffer. Serial DNA dilutions were quantified with a NanoDrop ND 1000 Spectrophotometer (NanoDrop Technologies).

Standard PCR was performed in a Veriti 96-well Thermal Cycler (Applied Biosystems) under the following conditions: template denaturation at 95°C for 1 min followed by 30 cycles of denaturation at 94°C for 30 s, annealing at 62.5°C for 30 s, extension at 72°C for 60 s, and a final extension of 5 min at 72°C. Primers used are listed in Supplementary Table 1.

### RNA extraction and PCR

Total RNA was extracted from tissues and cell cultures using Trizol (Sigma-Aldrich), followed by treatment with RNase-free DNAse (Roche Applied Science). cDNA was synthesized from 500 ng of total RNA using the Superscript III First Strand cDNA Synthesis Kit (Life Technologies). Specific mRNAs were quantified by qRT-PCR using an ABI Prism 7900 HT Detection System (Applied Biosystems) and TaqMan detection. PCR was performed in 96-well plate microtest plates with TaqMan master mix (Thermo Fisher) for 40 cycles. In all experiments, mRNA amounts were normalized to the total amount of cDNA by using amplification signals for hGUSB. Each sample was determined in triplicate, and at least three independent samples were analysed for each experimental condition.

### Western Blot

Proteins were extracted by standard procedures as previously described (Rodriguez-Perales et al. 2015) in the presence of Complete Protease Inhibitor Cocktail Tables (Roche Applied Science). Proteins were transferred to PVDF using TransFi (Invitrogen; Life Technologies), and membranes were probed for FOXP2 or MDFIC with monoclonal mouse anti-human FOXP2 or MDFIC antibodies (1/1000 or 1/500; BD Pharmigen) or for GAPDH (AbCam), with antibodies diluted 1/2500 in PBS/0.1% Tween-20 (PBS-T). Secondary antibodies were HRP-conjugated with goat anti-mouse IgG (1/1000) and goat anti-Rabbit (1/500; Dako, Barcelona, Spain), and blots were developed with ECL (GE Healthcare).

### Statistical Analysis

Data from three or more independent experiments were analysed by two-tailed unpaired t-test. NS, non-significant; * p<0.05; ** p<0.01; *** p<0.001; and **** p<0.0001.

## RESULTS

### In silico search of enhancer regions

We first hypothesised that the breakpoint in 7q31.1 (114,888,284 hg38) affected the expression of *FOXP2* by physically separating some cis-acting distant element with an enhancer role. Accordingly, we searched in silico for putative enhancers in the intergenic region between *FOXP2* and *MDIFC* looking for the following hallmarks: DNase I hypersensitive sites, presence of histones with specific post-translational modifications (specifically histone H3, lysine 4 monomethylation (H3K4me1) and H3 lysine 27 acetylation (H3K27ac)), and ChIP-seq data provided by ENCODE of regions recruiting co-activators and co-repressors as revealed by chromatin immunoprecipitation followed by deep sequencing. We found two putative enhancers located at 120kb and 203.5kb downstream the stop codon of *FOXP2*, respectively (Figure 1A and 1B). These putative enhancers (referred as FOXP2-E^proximal^ and FOXP2-E^distal^) span 6264bp (chr7:114,817,431-114,823,694 hg38) and 2300bp (chr7:114,900,989-114,903,302 hg38 equivalent to 114,541,370-114,542,201 hg19), respectively. FOXP2-E^distal^ is the one previously validated by luciferase assay by Becker et al. (2015); FOXP2-E^proximal^is a new putative regulatory element.

**Figure 1.**
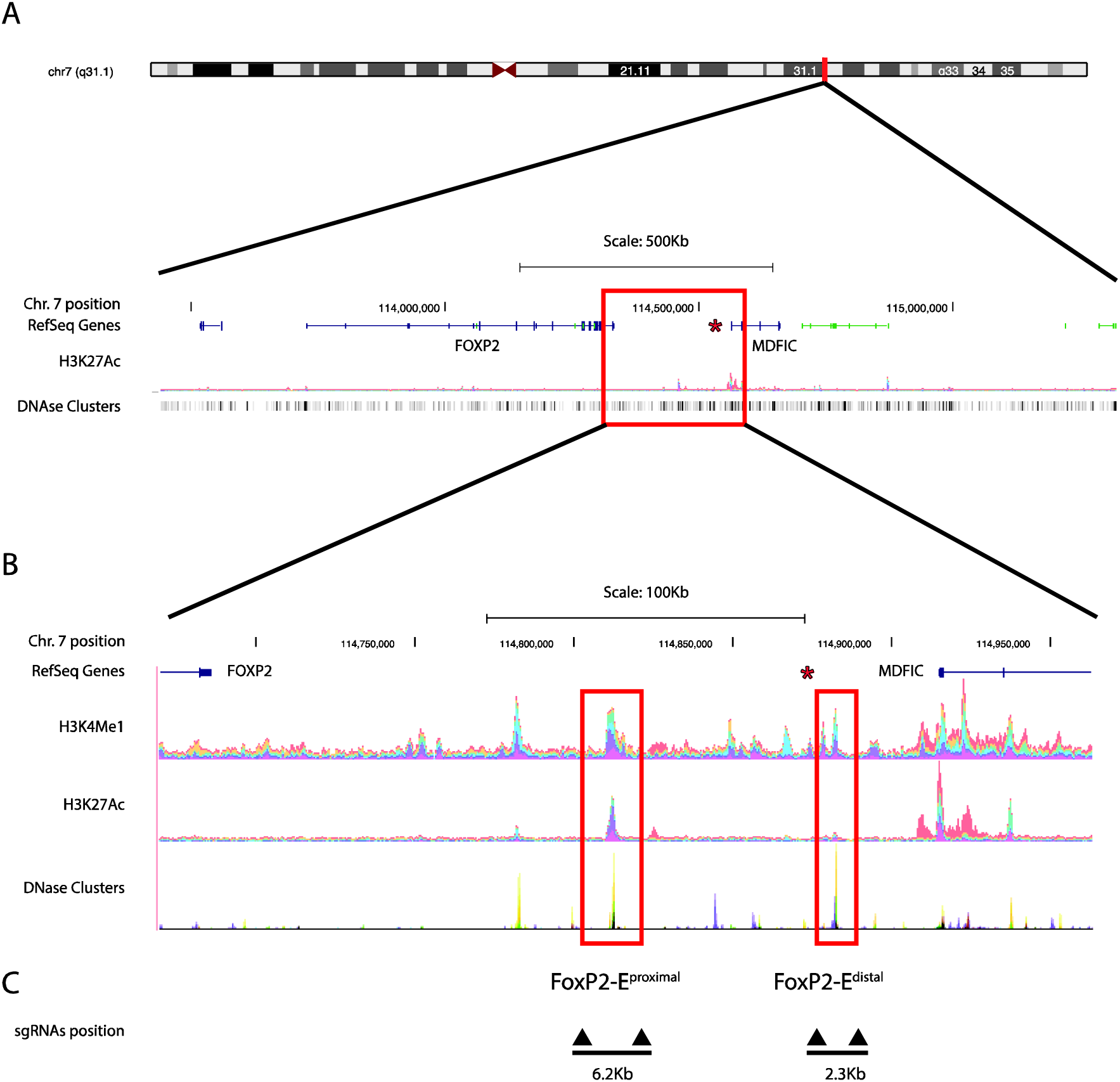
Identification of enhancer regions downstream *FOXP2* and upstream *MDFIC*. **A.** Genomic location of human *FOXP2* and *MDFIC* genes. Genes are depicted in blue. **B.** Detailed view of an Encode UCSC genome-browser snapshot showing bar graphs with a detailed representation of the locations of H3K4Me1 and H3K27Ac histone marks in human neurological cell lines. The squared regions in red show the locations of FOXP2-E^proximal^ and FOXP2-E^distal^. The red asterisk shows the position of the 7q breakpoint in the proband harbouring a genomic complex rearrangement and with severe expressive and receptive speech and language impairment. **C.** Schematic representation of the location of the four sgRNAs flanking the 6.2kb region including FOXP2-E^proximal^ and the 2.3kb region including FOXP2-E^distal^.

### CRISPR deletion of FOXP2-E^proximal^ and FOXP2-E^distal^

We then tested in vitro the functionality of FOXP2-E^proximal^ and FOXP2-E^distal^. Since both putative enhancers are located in an intergenic region, we aimed at characterizing that both of them are functional with respect to *FOXP2* or/and *MDFIC*. We relied on a CRISPR genome editing approach to delete the entire predicted sequence of each enhancer. Accordingly, we designed two couples of sgRNAs targeting the flanking regions of either FOXP2-E^proximal^ or FOXP2-E^distal^ (Figure 1C). Each sgRNA pair was cloned in the pLV-U6^#1^H1^#2^-C9G (Torres et al. 2014b) in order to couple the expression of the sgRNAs to the expression of Cas9 and a GFP reporter. Subsequently, we tested if the sgRNAs were able to induce the expected deletions. HEK293A cells were nucleofected with 2ug of pLV-U6^#1^H1^#2^-C9G plasmid targeting either FOXP2-E^proximal^ or FOXP2-E^distal^. After 72 h, the DNA was isolated and analyzed. After designing PCR oligos that span the deleted regions (Supplementary Table 1), PCR assays were performed. They revealed efficient targeted deletions of the 6.2kb or the 2.3kb regions containing the entire sequence of FOXP2-E^proximal^and FOXP2-E^distal^, respectively (data not shown).

### Neuronal cell lines defective in FOXP2-E^proximal^ or FOXP2-E^distal^

We next used CRISPR to delete FOXP2-E^proximal^or FOXP2-E^distal^ in two cell lines: SK-N-MC, a neuroblastomic cell line derived from the supra-orbital area, which expresses *FOXP2* constitutively (although it does not express *MDFIC* at the same level), and SH-SY5Y, a neuroblastomic cell line which expresses neither *FOXP2* nor *MDFIC*. The SH-SY5Y cell line was used as a negative control to confirm that changes produced by CRISPR-Cas9 were not due to unspecific effects. Cells were electroporated with 2ug of either pLV-U6^#1^H1^#2^-C9G- E^proximal^, pLV-U6^#1^H1^#2^-C9G-E^distal^, or with empty plasmids. After 72h the DNA was extracted and analysed. PCR and Sanger sequencing analyses confirmed the deletion of the 6.2kb or the 2.3kb fragment (Figures 2A and 2B). We then generated two clonal cell lines (one for each putative enhancer) by sorting GFP positive cells into 96-well plates for single cell colony expansion. We confirmed by PCR that each expanded cellular clone harbours a deletion containing either the FOXP2-E^proximal^or the FOXP2-E^distal^ regions (data not shown). These two cell lines were used for further expression analyses.

**Figure 2.**
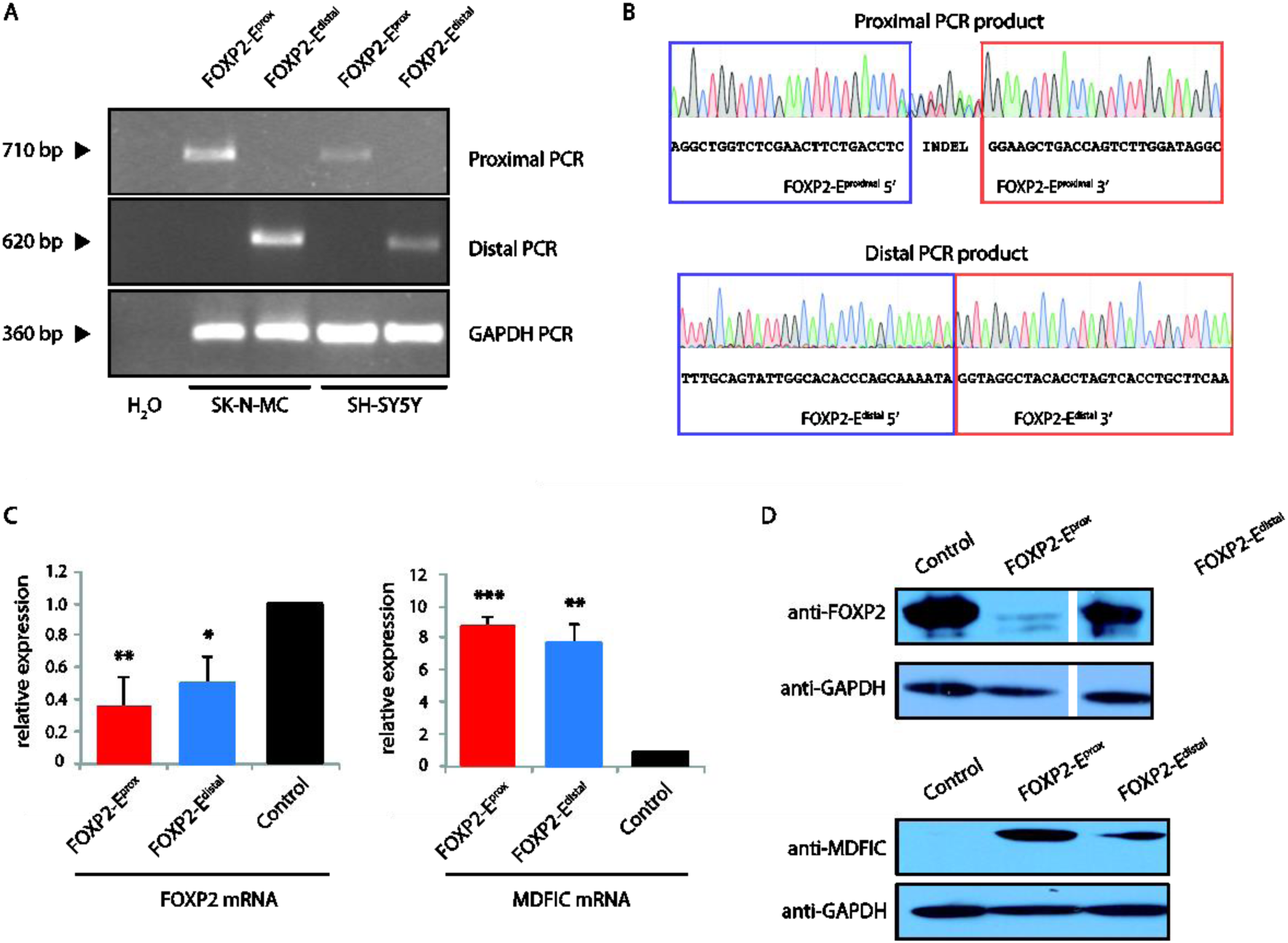
Molecular characterization of FOXP2-E^proximal^ and FOXP2-Edistal. **A.** PCR analysis. Two oligos flanking the deleted regions were used to amplify the genomic DNA from mutant SK-N-MC and SH-SY-5Y clones. GAPDH genomic oligos were used to amplify a positive control region. Black triangles show the size of the PCR products. **B.** Representative Sanger sequencing chromatogram showing the boundaries of the FOXP2-E^proximal^ and FOXP2-E^distal^ deleted regions in SK-N-MC cells. **C**. RT-qPCR analysis of SK-N-MC cells with FOXP2-E^proximal^ or FOXP2-E^distal^ deletions. Samples are normalized to the average *FOXP2* (left) or *MDFIC* (right) signal between three SK-N-MC wild type replicates. Levels of expression of *FOXP2* and *MDFIC* are represented by the fold change relative to that of parental control cell line, which were normalized to 1. **D.** Western blot analysis of cell lysates of SK-N-MC with FOXP2-E^proximal^ deleted, with FOXP2-E^distal^ deleted, and of control cells immunoblotted for FOXP2 (top) or MDFIC (bottom).

### *FOXP2* and *MDFIC* expression analyses

We next aimed to characterize in more detail the functionality of FOXP2-E^proximal^and FOXP2-E^distal^. We used RT-qPCR to determine the amount of *FOXP2* mRNA in the SK-N-MC cells transduced with either pLV-U6^#1^H1^#2^-C9G-E^proximal^, or pLV-U6^#1^H1^#2^-C9G-E^distal^, or an empty plasmid as a control. The expression of *FOXP2* was significantly reduced (2.9 fold change) compared to that of the control when FOXP2-E^proximal^was deleted (Figure 2C, left). Likewise, *FOXP2* expression was decreased (2 fold change) when FOXP2-E^distal^was deleted (Figure 2C left). We then measured the levels of expression of *MDFIC* after deletion of each enhancer. As shown in Figure 2C right, the expression of *MDFIC* was significantly increased when either FOXP2-E^proximal^ or FOXP2-E^distal^ were deleted (8.6 and 7.5 fold change, respectively).

The experiment was replicated in SH-SY5Y cells, but no change of expression of *FOXP2* or *MDFIC* was detected (data not shown). We next analysed by Western blot the amount of FOXP2 and MDFIC proteins in the SK-N-MC cell clones harbouring the FOXP2-E^proximal^or FOXP2-E^distal^ deletions or control cells transduced with an empty plasmid (Figure 2D). In line with our mRNA data, the deletion of FOXP2-E^proximal^or FOXP2-E^distal^ was found to reduce the level of FOXP2 (Figure 2D top) and to increase the level of MDFIC (Figure 2D bottom).

## DISCUSSION

In this paper we have characterised in detail the role of two functional regulatory elements located downstream *FOXP2*. One of these enhancers, FOXP2-E^distal^, had been previously found to be functional in a luciferase assay (Becker et al. 2015). We have been able to prove that if deleted, *FOXP2* becomes downregulated and the levels of FOXP2 protein are reduced in the SK-N-MC neuroblastomic cell-line. We have further proved that it also affects the adjacent gene, *MDFIC*. Its deletion upregulates *MDFIC* and increases the levels of MDFIC protein. The second enhancer, FOXP2-E^proximal^ was previously unknown. We have found that its deletion also downregulates *FOXP2* and upregulates *MDFIC*. These findings are coherent with previous studies reporting pairs of genes being governed by the same regulatory regions (Gould et al. 1997; Tsujimura et al. 2010), which in some cases have proven to regulate recruitment of RNA polymerase II to promoters of both genes (Collins et al. 2012).

*FOXP2* is a well-known gene, important for speech and language (Fisher and Scharff 2009, Graham and Fisher 2013). Less is known about the role of *MDFIC* in cognitive development and disease. There is evidence that the gene might be associated to language development and cognitive impairment (DECIPHER patients 262086, 292652, and 301696). *MDFIC* is a MyoD family inhibitor domain containing protein that acts as an activator or repressor of transcription (Thebault et al. 2000, Gautier et al. 2005). Similarly to FOXP2, it interacts with LEF1, as part of beta-catenin regulation (Kusano et al. 2002). In chicken MDFIC is targeted by miR-130a (Han et al. 2016), known to regulate neurite outgrowth and dendritic spine density by targeting *MeCP2*, the main candidate for Rett syndrome, a condition involving language deficits (Zhang et al. 2016). *MDFIC* is highly expressed in the cerebellum during human embryonic development and in the thalamus after birth (Human Brain Transcriptome http://hbatlas.org/). These two brain regions, interacting in a dopaminergic cortico-striato-thalamic loop, seem to play an important role in timing sensorimotor control, needed for auditory-motor language processing (Alcock et al. 2000, Vargha-Khadem et al. 2005). The cerebellum and thalamus of those bearing the R553H mutation in *FOXP2,* associated to speech and language impairment, exhibit volumetric changes of their grey matter (Watkins et al. 2002a). This suggests that the modulation of the size of neural populations in particular regions in the loop may impact in sensorimotor performance.

Our findings in a human neuronal cell line give support to the view that the breakpoint in our proband, which separated each of these two intergenic enhancers from each other and consequently from one of the two genes they regulate, may have altered the expression levels of both *FOXP2* and *MDFIC* contributing to the observed speech and language deficits (Moralli et al 2015). Whereas *MDFIC* remained in chromosome 7q with, predictably, FOXP2-E ^distal^, *FOXP2* was rearranged to chromosome 11p with, predictably, FOXP2-E^proximal^. Accordingly, we expect the expression of *FOXP2* to be down-regulated and the expression of *MDFIC* to be up-regulated. The knockdown of *FoxP2* in zebra finch results in a shorter window for song learning and in less accurate and shorter song imitation (Haesler et al., 2007). This is coherent with the inability that those carrying the R553H mutation of *FOXP2* show in repeating words and pseudowords (Watkins et al., 2002b). Further confirmation of our hypothesis would need to be supported in a specific neuronal brain cell line grown from stem cells of the proband, as well as in songbirds in which these enhancers have been deleted.

We expect that our findings also contribute to a better understanding of the role that this region may have played in the evolution of language. Differences in the expression levels of both *FOXP2* and *MDFIC* are expected between extinct hominins and modern humans, plausibly accounting for some of the presumed differences in their language abilities. Neanderthals bear the ancestral allele of a binding site for POU3F2 within intron 8 of *FOXP2*, which is more efficient in activating transcription (Maricic et al. 2013). Accordingly, higher levels of FOXP2 are expected in this hominin species. Likewise, the *MDFIC* locus is among the top five percent S score regions in modern humans (Green et al. 2010, table S37). Finally, both genes are functionally related to *RUNX2*, which encodes an osteogenic factor that controls the closure of cranial sutures and several aspects of brain growth, and that has been related to the changes that brought about our more globular brain (case) and our species-specific mode of cognition, including language (Boeckx and Benítez-Burraco 2014; Benítez-Burraco and Boeckx 2015). Further confirmation of this hypothesis would need to be supported by the analysis of the enhancers’ sequences in extinct hominins and by mimicking the attested changes (if any) in a human cell line.

Ii is expected that our study, together with new available data about seed sequences of miRs in the 3’UTR region of *FOXP2* (Clovis et al. 2012; Shi et al. 2013; Fu et al. 2014a; Cuiffo et al. 2014), contributes to a deeper understanding of how *FOXP2* is regulated. Since we have found that both adjacent genes, *FOXP2* and *MDFIC*, are sharing a regulatory region downstream the 3’ end of *FOXP2* which modulates their expression, it is expected that these results will contribute as well to a better understanding of how each gene participates in the development of subcellular and sensorimotor neural function underlying language.

## ACKNOWLEDGMENTS

This project was supported by funds from The University of Oxford John Fell OUP Research Grant [121/435] awarded to its Principal Investigator Paloma Garcia-Bellido. This study was supported in part by funds from the Spanish National Research and Development Plan, Instituto de Salud Carlos III, and FEDER (FIS project no. PI14/01884 to Sandra Rodriguez-Perales), and in part by funds from the Spanish Ministry of Economy and Competitiveness (grant numbers FFI2014-61888-EXP and FFI-2013-43823-P to Antonio Benítez-Burraco). The funders had no role in the study design, data collection and analysis, decision to publish, or preparation of the manuscript. We thank Daniela Moralli, Dianne Newbury and Sonja C. Vernes for their helpful comments.

## ETHICS

Ethics approval for this research was granted by the University of Oxford [MSD-IDREC-C1-2012-95, SSD/CUREC2/09-23].

## AUTHORS CONTRIBUTION

P G-B conceived the project, participated in the design and coordination of experiments, analysed results, and wrote the paper.

A B-B analysed results and wrote the paper.

R T-R did the experiments, analysed results, and wrote the paper.

M M-L did the experiments.

S R-P participated in the design, coordinated experiments, analysed results, and wrote the paper.

All authors read and approved the final manuscript

---

**Supplementary Table 1.** Oligonucleotide and sgRNA sequences used in this study.

## sgRNA sequences (IDT gblocks)

**sgRNA_FOXP2_Edistal_1_5:**

ccataACGCGTTGTACACGAACGCTGACGTCATCAACCCGCTCCAAGGAATCGCGGGCCCAGTGTCACTAGGCGGGAACACCCAGCGCGCGTGCGCCCTGGCAGGAAGATGGCTGTGAGGGACAGGGGAGTGGCGCCCTGCAATATTTGCATGTCGCTATGTGTTCTGGGAAATCACCATAAACGTGAAATGTCTTTGGATTTGGGAATCTTATAAGTTCTGTATGAGACCACTCTTTCCC**G**<u>cacacccagcaaaatacat</u>GTTTTAGAGCTATGCTGGAAACAGCATAGCAAGTTAAAATAAGGCTAGTCCGTTATCAACTTGAAAAAGTGGCACCGAGTCGGTGCTTTTTTACTAGTcgcta

## sgRNA_FOXP2_Edistal_2_5

ccataACGCGTTGTACACGAACGCTGACGTCATCAACCCGCTCCAAGGAATCGCGGGCCCAGTGTCACTAGGCGGGAACACCCAGCGCGCGTGCGCCCTGGCAGGAAGATGGCTGTGAGGGACAGGGGAGTGGCGCCCTGCAATATTTGCATGTCGCTATGTGTTCTGGGAAATCACCATAAACGTGAAATGTCTTTGGATTTGGGAATCTTATAAGTTCTGTATGAGACCACTCTTTCCC**G**<u>gcaaggtatattctctgag</u>GTTTTAGAGCTATGCTGGAAACAGCATAGCAAGTTAAAATAAGGCTAGTCCGTTATCAACTTGAAAAAGTGGCACCGAGTCGGTGCTTTTTTACTAGTcgcta

## sgRNA_FOXP2_Edistal_1_3

ccataCAATTGGGGCAGGAAGAGGGCCTATTTCCCATGATTCCTTCATATTTGCATATACGATACAAGGCTGTTAGAGAGATAATTAGAATTAATTTGACTGTAAACACAAAGATATTAGTACAAAATACGTGACGTAGAAAGTAATAATTTCTTGGGTAGTTTGCAGTTTTAAAATTATGTTTTAAAATGGACTATCATGTACACTTACCGTAACTTGAAAGTATTTCGATTTCTTGGCTTTATATATCTTGTGGAAAGGACGAGGTACC**G**<u>atctactcttctttagggt</u>GTTTTAGAGCTATGCTGGAAACAGCATAGCAAGTTAAAATAAGGCTAGTCCGTTATCAACTTGAAAAAGTGGCACCGAGTCGGTGCTTTTTTACGCGTACTAGTcgcta

## sgRNA_FOXP2_Edistal_2_3

ccataCAATTGGGGCAGGAAGAGGGCCTATTTCCCATGATTCCTTCATATTTGCATATACGATACAAGGCTGTTAGAGAGATAATTAGAATTAATTTGACTGTAAACACAAAGATATTAGTACAAAATACGTGACGTAGAAAGTAATAATTTCTTGGGTAGTTTGCAGTTTTAAAATTATGTTTTAAAATGGACTATCATGTACACTTACCGTAACTTGAAAGTATTTCGATTTCTTGGCTTTATATATCTTGTGGAAAGGACGAGGTACC**G**<u>gaagagtagatcgcatgag</u>GTTTTAGAGCTATGCTGGAAACAGCATAGCAAGTTAAAATAAGGCTAGTCCGTTATCAACTTGAAAAAGTGGCACCGAGTCGGTGCTTTTTTACGCGTACTAGTcgcta

## sgRNA_FOXP2_Eproximal_1_5

ccataACGCGTTGTACACGAACGCTGACGTCATCAACCCGCTCCAAGGAATCGCGGGCCCAGTGTCACTAGGCGGGAACACCCAGCGCGCGTGCGCCCTGGCAGGAAGATGGCTGTGAGGGACAGGGGAGTGGCGCCCTGCAATATTTGCATGTCGCTATGTGTTCTGGGAAATCACCATAAACGTGAAATGTCTTTGGATTTGGGAATCTTATAAGTTCTGTATGAGACCACTCTTTCCC**G**<u>gtgatctcagctactcggg</u>GTTTTAGAGCTATGCTGGAAACAGCATAGCAAGTTAAAATAAGGCTAGTCCGTTATCAACTTGAAAAAGTGGCACCGAGTCGGTGCTTTTTTACTAGTcgcta

## sgRNA_FOXP2_Eproximal_2_5

ccataACGCGTTGTACACGAACGCTGACGTCATCAACCCGCTCCAAGGAATCGCGGGCCCAGTGTCACTAGGCGGGAACACCCAGCGCGCGTGCGCCCTGGCAGGAAGATGGCTGTGAGGGACAGGGGAGTGGCGCCCTGCAATATTTGCATGTCGCTATGTGTTCTGGGAAATCACCATAAACGTGAAATGTCTTTGGATTTGGGAATCTTATAAGTTCTGTATGAGACCACTCTTTCCC**G**<u>ctcgaacttctgacctcag</u>GTTTTAGAGCTATGCTGGAAACAGCATAGCAAGTTAAAATAAGGCTAGTCCGTTATCAACTTGAAAAAGTGGCACCGAGTCGGTGCTTTTTTACTAGTcgcta

## sgRNA_FOXP2_Eproximal_1_3

ccataCAATTGGGGCAGGAAGAGGGCCTATTTCCCATGATTCCTTCATATTTGCATATACGATACAAGGCTGTTAGAGAGATAATTAGAATTAATTTGACTGTAAACACAAAGATATTAGTACAAAATACGTGACGTAGAAAGTAATAATTTCTTGGGTAGTTTGCAGTTTTAAAATTATGTTTTAAAATGGACTATCATGTACACTTACCGTAACTTGAAAGTATTTCGATTTCTTGGCTTTATATATCTTGTGGAAAGGACGAGGTACC**G**<u>ctgtaataagatagcaggg</u>GTTTTAGAGCTATGCTGGAAACAGCATAGCAAGTTAAAATAAGGCTAGTCCGTTATCAACTTGAAAAAGTGGCACCGAGTCGGTGCTTTTTTACGCGTACTAGTcgcta

## sgRNA_FOXP2_Eproximal_2_3

ccataCAATTGGGGCAGGAAGAGGGCCTATTTCCCATGATTCCTTCATATTTGCATATACGATACAAGGCTGTTAGAGAGATAATTAGAATTAATTTGACTGTAAACACAAAGATATTAGTACAAAATACGTGACGTAGAAAGTAATAATTTCTTGGGTAGTTTGCAGTTTTAAAATTATGTTTTAAAATGGACTATCATGTACACTTACCGTAACTTGAAAGTATTTCGATTTCTTGGCTTTATATATCTTGTGGAAAGGACGAGGTACC**G**<u>tatggctgccacattccgt</u>GTTTTAGAGCTATGCTGGAAACAGCATAGCAAGTTAAAATAAGGCTAGTCCGTTATCAACTTGAAAAAGTGGCACCGAGTCGGTGCTTTTTTACGCGTACTAGTcgcta

## Primers qPCR

~~~
> qFoxP2_Fw
GCAGCAGAGATGGAAGATCA
> qFoxP2_Rv
AGTTGTCTTGCTGCCTGGAG
cDNA amplicon size:103            Estimated genomic amplicon size:108040
> qMDFIC_Fw
GTCCATTTGGGGAAATCCTT
> qMDFIC_Rv
CATTGCTCAGACCTGTGTGG
cDNA amplicon size:140            Estimated genomic amplicon size:37248
~~~

## Primers Surveyor & deletion detection

~~~
sMDFIC Fw2
TGATCTCAGTGCAGGCAAA
sMDFIC Rv2
GTTGGACTAAGGTGCCAGTT

                                  2314pb (deletion FOXP2 Distal)

sMDFIC Fw
TACTGTTTCATGGATGCTGACT
sMDFIC Rv
CCTTTGGCCACAGACTGAA

sFOXP2 Fw
GGGATAGCACTGGGAGAAATAC
sFOXP2 Rv
GCGGTGGCTCATTTCTGTA

                                  6264 pb (deletion FOXP2 Proximal)

sFOXP2 Fw2
TTCTGCACCTTGGGTTAGG
sFOXP2 Rv2
AGGGTTGATTGATTGCCAGAG
~~~

